# Transcriptional regulation of *SLIT2* expression in pancreatic cancer cell lines

**DOI:** 10.1101/2020.09.29.319129

**Authors:** Brenna A. Rheinheimer, Lukas Vrba, Bernard W Futscher, Ronald L Heimark

## Abstract

**Background:** SLIT2 has been shown to serve as a tumor suppressor in breast, lung, colon, and liver cancers. Additionally, expression of SLIT2 has been shown to be epigenetically regulated in prostate cancer. Therefore, we sought to determine transcriptional regulation of SLIT2 in pancreatic ductal adenocarcinoma.

**Methods:** RNA expression of SLIT2, SLIT3, and ROBO1 was examined in a panel of pancreatic ductal adenocarcinoma cell lines while protein expression of ROBO1 and SLIT2 was examined in tumor tissue. Methylation of the SLIT2 promoter was determined using Sequenom while histone modifications were queried by chromatin immunoprecipitation. Reexpression of SLIT2 was tested by treatment with 5-aza-2’deoxycytidine and Trichostatin A.

**Results:** Pancreatic cancer cell lines fall into three distinct groups based on SLIT2 and ROBO1 expression. The SLIT2 promoter is methylated in pancreatic ductal adenocarcinoma and SLIT2 expression is dependent on the level of methylation at specific CpG sites. Treatment with 5-aza-2’deoxycytidine (but not Trichostatin A) led to SLIT2 reexpression. The SLIT2 promoter is bivalent in pancreatic ductal adenocarcinoma and histone marks around the transcriptional start site are responsible for transcription.

**Conclusions:** Loss of SLIT2 expression modulated by epigenetic silencing may play a role in pancreatic ductal adenocarcinoma progression.

## Introduction

To date, there are no effective treatment options for patients with metastatic pancreatic ductal adenocarcinoma. We lack understanding of 1) genes regulating pancreatic cancer invasion and 2) the mechanisms behind which these unknown genes operate to promote invasion of this disease. My studies show that pancreatic ductal adenocarcinoma cell lines can be classified into three distinct subtypes based on *ROBO1* and *SLIT2* mRNA expression. Transcription of intronic mir-218-1 is uncoupled from *SLIT2* mRNA expression and is transcriptionally controlled through a candidate alternative promoter located within intron 4 of the *SLIT2* gene. Suppression of pancreatic cancer invasion through this *SLIT2*/mir-218-1 signaling axis occurs via a novel miR-218 target, ARF6, and inhibition of extracellular matrix degradation by invadopodia.

Previous studies on mammary duct branching morphogenesis, similar to the proliferation of ductal structures during pancreatic ductal adenocarcinoma progression, identified a type of epithelial collective cell migration that does not include cellular extensions or protrusions^223^. Ewald et al. showed that cells within extending mammary ducts dynamically rearrange, but remain adherent. During this collective migration, the tissue continues to have distinct luminal- and basement membrane-contacting surfaces, yet the individual cells within the multilayered region are incompletely polarized suggesting that collective epithelial cell migration drives mammary duct branching morphogenesis. Myoepithelial cells also play an important role in mammary development. In the same study, Ewald et al. also discovered a close relationship between myoepithelial cell localization, myoepithelial cell motility and the resulting pattern of branching morphogenesis suggesting an important role for myoepithelial cells as cellular regulators of tissue structure. Axon guidance molecules have been shown to play an important part in mammary branching morphogenesis as well. SLIT/ROBO1 signaling has been shown to inhibit lateral branch formation by controlling proliferation of the basal cell layer^219^ suggesting a potential role for SLIT/ROBO1 signaling in collective cell migration. Considering the pancreas also contains a ductal structure that proliferates during pancreatic ductal adenocarcinoma formation and progression, understanding the role of axon guidance molecules in pancreatic cancer would open up new mechanisms through which pancreatic ductal adenocarcinoma cells migrate and invade to form metastases. Similar studies on SLIT/ROBO signaling have also been shown for growth and invasion of blood vessels during angiogenesis^214,224,225^.

The axon guidance ligand *SLIT2* is an attractive molecule to study in this context for several reasons. During mouse mammary gland duct development, loss of *Slit* gene expression leads to the formation of hyperplastic, disorganized ducts with a coordinate increase in desmoplastic stroma surrounding the perturbed ductal structures^209^ suggesting a role for Slits in cancer formation. *SLIT2* has also been shown to have tumor suppressor qualities in epithelial cancers including decreased colony formation^205^, induction of apoptosis^206^, tumor growth *in vivo*^209^, cell migration and invasion^207^, and metastasis^211^. Genomic aberrations of SLIT2 have also been shown in early-stage pancreatic ductal adenocarcinoma lesions with 7% of patients showing loss of *SLIT2* and 3% of patients containing a mutation in *SLIT2*^220^. Therefore, determining the role of *SLIT2* in pancreatic cancer, specifically the mechanism behind *SLIT2*-mediated invasion, would vastly improve the understanding of the complexity of pancreatic ductal adenocarcinoma invasion and perhaps provide a biomarker to stratify high-risk patients or a molecular target for novel treatment therapies.

A CpG island is defined as a segment of DNA greater than 200 bp with a GC content greater than 50% and an observed-to-expected CpG ratio of greater than 0.6^226^. Using the EpiDesigner website from Sequenom, *SLIT2* was found to contain a very large CpG island that is 3592 bp long, has a GC content of 66%, and an observed-to-expected CpG ratio of 0.767. The CpG island in the *SLIT2* promoter starts at −2021 bp upstream of the transcriptional start site, runs through the first coding exon, and ends within the first intron at +1571 bp downstream of the transcriptional start site. It has already been shown that HDAC5, a class II histone deacetylase, binds to the *SLIT2* promoter in human umbilical vein endothelial cells to suppress angiogenesis by inhibiting capillary-like sprouting of endothelial cells^225^. The basic helix-loop-helix transcription factor NeuroD1 was also found to bind to the first and second E-box of the *Slit2* promoter in *MYCN* transgenic mice resulting in suppressed *Slit2* expression and increased neuroblastoma cell motility^227^. This signifies that transcriptional regulation of *SLIT2* is a complex process likely to depend on a number of factors and cellular context.

Bivalent promoters contain chromatin enrichment of methylation on H3K4 and H3K27me3 simultaneously and are considered to poise expression of genes allowing for timely activation of gene expression while maintaining transcriptional repression in the absence of activating signals^228^. Specific DNA sequence elements, DNA methylation status, specific histone modifications, and transcription factors have all been implicated in the generation of bivalent domains^229^. CpG islands have also been shown to correlate with and play a role in establishing bivalent domains since essentially all H3K4me3 sites map to CpG islands^230^. CpG islands also play a role in establishing and maintaining H3K27me3 at bivalent domains, though not all CpG islands are marked with enrichment of H3K27me3. Approximately 97% of all promoter-associated EZH2 recruitment sites correspond to CpG islands^231^. Transcription of bivalent domains depends on a number of factors including the presence of EZH2, the presence of transcriptional repressors, and the presence of transcriptional activators. Poised promoters can be defined as a bivalent promoter that also contains chromatin enrichment for a poised form of RNA polymerase II that is phosphorylated at Serine 5^232^. Examples of tumor suppressor genes that contain bivalent promoters include *CDH1, CDKN2A, RASSF1A, GATA4*, and *GATA5^233^* suggesting that this mechanism of transcriptional regulation may be widely utilized for tumor suppressor genes throughout the genome. Bivalent promoters of *CDH1, CDKN2A*, and *RASSF1A* have been seen in pancreatic cancers as well^81,234^. Since *SLIT2* gene expression has previously been shown to be affected by both DNA CpG methylation^205–207^ and chromatin enrichment of H3K27me3^208^, I proposed that *SLIT2* contains a bivalent domain and that its expression depends on specific factors present in the nucleus during transcription.

Based on findings within the literature, the first aim of this dissertation was to identify the mechanism through which *SLIT2* expression is lost in pancreatic ductal adenocarcinoma. *The hypothesis driving this aim is that SLIT2 expression is lost in pancreatic cancer through both DNA methylation at the gene promoter and chromatin remodeling at the transcriptional start site*. Quantitative reverse transcription PCR was used to measure the expression of several members of the Slit family of ligands and the Robo family of receptors in a panel of pancreatic cancer cell lines. Sequenom was utilized to quantitatively elucidate the DNA methylation landscape of the *SLIT2* promoter. Treatment of pancreatic cancer cell lines with the demethylating agent 5-aza-2’deoxycytidine was performed to establish if *SLIT2* gene expression could be turned back on in cells with no detectable *SLIT2* mRNA expression. Finally, chromatin immunoprecipitation was used to measure the chromatin enrichment of histone modifications present at the *SLIT2* promoter and transcriptional start site.

## Materials and methods

### Cell culture and treatment

Immortalized human pancreatic ductal epithelial cells were obtained from the Ming-Sound Tsao lab at the University of Toronto, Toronto, Ontario, Canada and maintained in keratinocyte serum-free media supplemented with 0.05 mg/ml of bovine pituitary extract and 0.02 μg/ml EGF recombinant human protein (Life Technologies, Benicia, CA) in a 37°C, 5% CO_2_ atmosphere with constant humidity. BxPC-3, Su.86.86, HPAF-II, and Hs 766T were obtained from the American Tissue Culture Collection (Manassas, VA) and maintained in Roswell Park Memorial Institute (RPMI) 1640 medium with 10% heat-inactivated fetal bovine serum (Life Technologies, Benicia, CA) and penicillin/streptomycin (Life Technologies, Benicia, CA) in a 37°C, 5% CO_2_ atmosphere with constant humidity. MIA PaCa-2 and PANC-1 were obtained from the American Tissue Culture Collection (Manassas, VA) and maintained in Dulbecco’s Modified Eagle Medium (DMEM) with 10% heat-inactivated fetal bovine serum and penicillin/streptomycin (Life Technologies, Benicia, CA) in a 37°C, 5% CO_2_ atmosphere with constant humidity. Capan-1 and Capan-2 were obtained from the American Tissue Culture Collection (Manassas, VA) and maintained in McCoy’s 5A medium (Sigma Aldrich, St. Louis, MO) with 10% heat-inactivated fetal bovine serum and penicillin/streptomycin (Life Technologies, Benicia, CA) in a 37°C, 5% CO_2_ atmosphere with constant humidity. The genotype of each pancreatic cancer cell line is shown in Table 2.1.

**Table 2.1:** Genotype of pancreatic cancer cell lines. Genetic alterations in several genes known to be involved with pancreatic cancer formation and progression. HD = homozygous deletion and -- = no genetic alteration.

| **Cell Line** | **KRAS** | **p16** | **p53** | **SMAD4** |
| --- | --- | --- | --- | --- |
| BxPC-3 | Wild-type | HD | Y220C | HD |
| Su.86.86 | G12D | HD | G245S | -- |
| Capan-1 | G12V | HD | A159V | Early stop |
| Capan-2 | G12V | Insertion | -- | -- |
| HPAF-II | G12D | In-frame deletion | P151S | -- |
| MIA PaCa-2 | G12C | HD | R248W | -- |
| PANC-1 | G12D | HD | R273H | -- |
| Hs 766T | Q61H | Splice mutation | Rearrangement | HD |

For treatment, BxPC-3, Su.86.86, Capan-1, Capan-2, PANC-1, and Hs 766T were plated at a density of 2×10^4^ cells/cm^2^ and were treated for six days with 100 μM 5-aza-2’deoxycytidine (Sigma Aldrich, St. Louis, MO) or for 24 hours with 50 nM Trichostatin A (Sigma Aldrich, St. Louis, MO). For the combination treatment with both 5-aza-2’deoxycytidine and Trichostatin A, cells were plated at a density of 2×10^4^ cells/cm^2^ and treated for six days with 100 μM of 5-aza-2’deoxycytidine with 50 nM Trichostatin A added 24 hours before the cells were collected in Trizol. Total RNA was isolated and reverse transcribed into cDNA and gene expression analyzed by quantitative reverse transcription PCR (qPCR). Cell lines were treated in duplicate and each RNA sample was analyzed by qPCR in triplicate.

### Nucleic acid isolation and quantitative reverse-transcription PCR (qPCR)

Total RNA was extracted using Trizol (Life Technologies, Benicia, CA) and quantified by absorption measurements at 260 nm. 1 μg of RNA was reverse transcribed using 1 μg/ml random primers and SuperScript II reverse transcriptase (Life Technologies, Benicia, CA). Intron-spanning oligo primers to *ROBO1, ROBO2, SLIT2, SLIT3, CDH1, ZEB1, DNMT1, DNMT3a, DNMT3b, EZH2* and *18s* rRNA (Table 2.2) were designed using the Roche Universal Probe Library assay design center. Each primer set was designed to a specific Roche Universal Probe Library probe containing a FAM quencher. Primers were ordered from Integrated DNA Technologies (Coralville, IA) and qPCR was performed using Quanta PerfeCTa Supermix, Low Rox (Quanta BioScience, Gaithersburg, MD) with the corresponding Roche Universal Probe Library probe on an ABI Prism 7500 Sequence Detection System (Life Technologies, Benicia, CA). Differences in expression between cancer cell lines and HPDE were determined using the comparative Ct method described in the ABI user manual relative to *18s* for *ROBO1, ROBO2, SLIT2, SLIT3, CDH1, ZEB1, DNMT1, DNMT3a, DNMT3b* and *EZH2*. Total RNA was isolated from each cell line in duplicate and each RNA sample was processed by qPCR in triplicate.

**Table 2.2:**
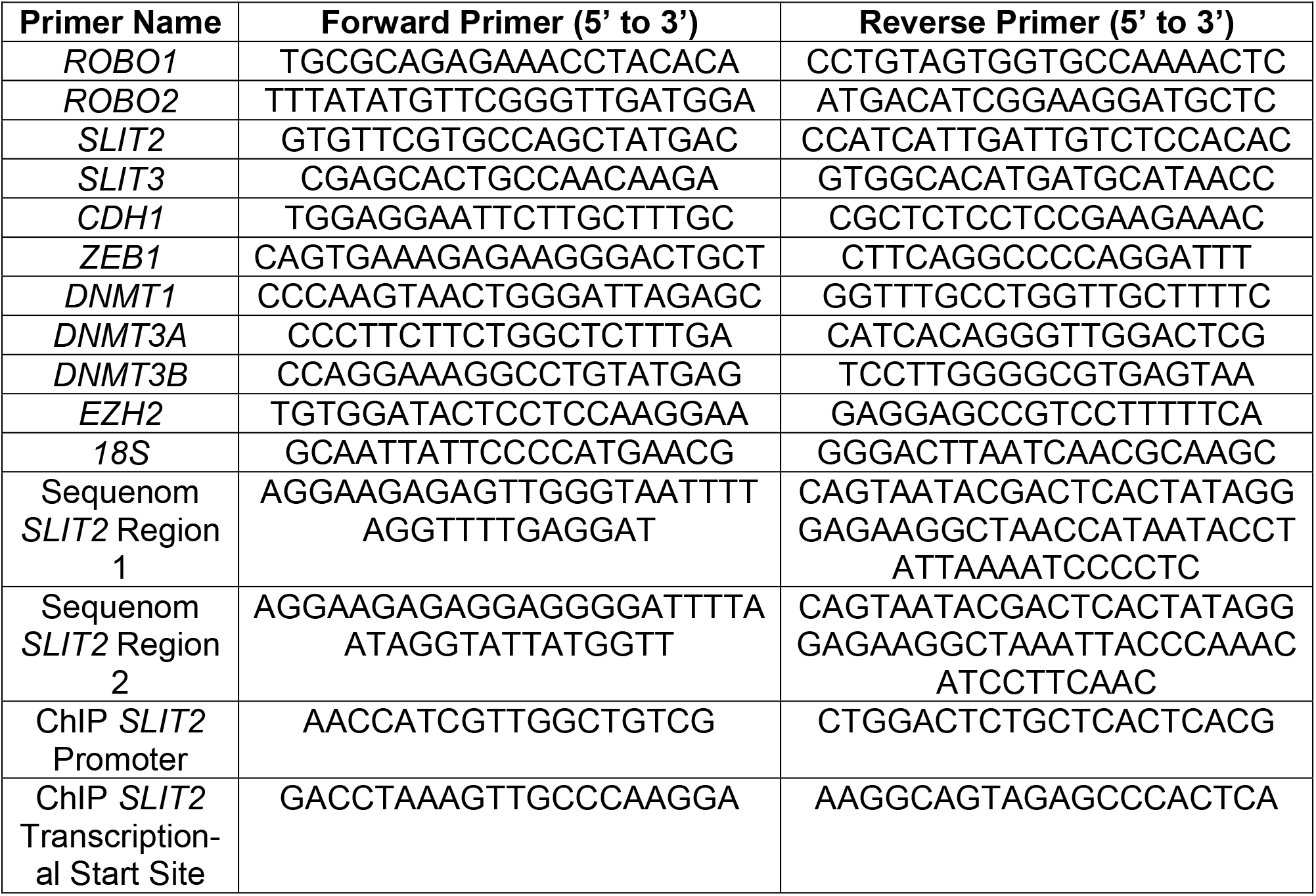
Primer sequences used in Chapter 2. Primer sequences used to determine gene expression, DNA CpG methylation via Sequenom analysis, and chromatin enrichment of histone modifications in Chapter 2. All primers are written from 5’ to 3’.

### Pancreatic clinical specimens and immunohistochemistry

Deidentified, formalin-fixed, paraffin-embedded archival tissue blocks were obtained from patients that had not received any preoperative chemotherapy or radiation therapy. All patients provided written consent and all phases of the research were approved by the Internal Review Board of the University of Arizona. Frozen tissue, as well as paired tumor and distant normal tissues were retrieved from the GI SPORE tissue bank. Pancreatic tissue sections were deparaffinized with Xylene and rehydrated. Antigen retrieval was carried out with 10 nM Citrate Buffer, pH 6 at 95-100°C for 30 minutes. Endogenous peroxidase was blocked with hydrogen peroxide in methanol and slides were incubated in blocking solution (3% normal donkey serum + 1% Tween20 in PBS) for 1 hour at room temperature. Primary antibody in blocking solution was added and incubated overnight at 4°C. The primary antibody towards ROBO1 (ab7279) was purchased from Abcam (Cambridge, United Kingdom) and the primary antibody towards SLIT2 (AB5701) was purchased from Millipore (Billerica, MA). Biotinylated secondary antibody was added and incubated for 30 minutes followed by incubation with Vectastain Elite avidin biotin complex for an additional 30 minutes. Slides were developed with 3,3-diaminobenzidine and counterstained with hematoxylin. The immunohistochemical stained slides were scanned at 40X and 200X.

### Sequenom analysis for DNA methylation

Genomic DNA was isolated from pancreatic cancer cell lines using the DNeasy Blood & Tissue Kit (Qiagen, Valencia, CA) and quantified spectrophotometrically. 500 ng of DNA was then sodium bisulfite converted using the Zymo Reseach Methylation Gold kit (Zymo Research, Irvine, CA). Primers to the *SLIT2* core promoter (from −611 bp to −317 bp and from −344 bp to −79 bp upstream of the *SLIT2* transcriptional start site) were designed using the EpiDesigner software (Sequenom, San Diego, CA). Primers used are listed in Table 2.2. Sodium bisulfite-treated DNA was amplified using a region-specific PCR that incorporated a T7 RNA polymerase sequence using part of the Sequenom MassARRAY PCR Reagents kit (Sequenom, San Diego, CA). The resulting PCR product was then *in vitro* transcribed and cleaved by RNase A using the MassCLEAVE T-only kit, spotted onto a Spectro CHIP array, and analyzed using the MassARRAY Compact System matrix-assisted laser desorption/ionization-time-of-flight mass spectrometer (Sequenom, San Diego, CA) at the University of Arizona Genetics Core. Each sodium bisulfite-treated DNA sample was processed in triplicate. Data was analyzed using the EpiTyper software (Sequenom, San Diego, CA). Data is presented as the average percent methylation within each fragment where yellow indicates 0% DNA methylation and dark blue indicates 100% DNA methylation. To calculate the average methylation level, the R-script “Analyze Seuqnom Function” weights each signal-to-noise *(SNR)* by the number of methylated CpG sites (the unmethylated peak is defined as *NOME*) represented in the peak using the following formula:

Average methylation of fragment with *n* CpG sites:

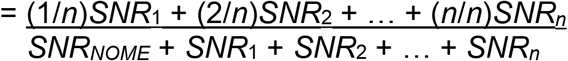

### Chromatin Immunoprecipitation (ChIP)

Cell lysates were collected and processed using the EZ-ChIP kit (Millipore, Billerica, MA). Briefly, cells were treated with 1% formaldehyde for 10 minutes to cross-link DNA and protein. Cells were then scraped from cell culture plates in CMF-PBS (Life Technologies, Billerica, CA) containing a protease inhibitor cocktail (Sigma Aldrich, St. Louis, MO). Resulting DNA-protein complexes were sonicated and subjected to gel electrophoresis to ensure proper sonication. Ten percent of the product was removed for later analysis as input DNA. The remaining portion was precleared using protein A Sepharose GL-4B beads (GE Healthcare, Piscataway, NJ) and incubated overnight with an antibody directed toward a specific histone modification. Antibodies against H3K4me2, H3K27me3, H4ac, and RNA polymerase II were purchased from Millipore (#07-030, #07-449, #06-866, and #05-623, respectively). After incubation, the bound DNA was immunoprecipitated, washed, and treated with 5 M NaCl to reverse DNA-protein cross-links, after which protein was digested with proteinase K (Life Technologies, Billerica, CA). Immunoprecipitated and input DNA samples were purified using the PCR Purification Kit (Qiagen, Valencia, CA) and were quantified spectrophotometrically. Each cell line was immunoprecipitated twice. Primers to the *SLIT2* core promoter (from −568 bp to −474 bp) and transcriptional start site (from −96 bp to −6 bp) were designed using the Roche Universal Probe Library assay design center. Primers used are listed in Table 2.2. Equal amounts of ChIP and input DNA were used for qPCR analysis. qPCR was performed using Power SYBR® Green Master Mix (Life Technologies, Benicia, CA) on an ABI Prism 7500 Sequence Detection System (Life Technologies, Benicia, CA). Differences in immunoprecipitation were determined using the fold enrichment method (Life Technologies, Benicia, CA). Each ChIP sample was analyzed by qPCR in triplicate.

## Results

### Axon guidance molecule expression in pancreatic cancer cell lines and primary pancreatic cancer tissue samples

Axon guidance molecules of the Slit and Robo family have recently been shown to be involved in pancreatic cancer neural invasion and metastasis. Göhrig et al. determined that restoration of SLIT2 expression in SLIT2-deficient pancreatic ductal adenocarcinoma cells inhibited their bidirectional chemoattraction to neural cells and impaired their unidirectional navigation along neural outgrowths. Conversely, silencing of ROBO1 stimulated the migration of SLIT2-containing pancreatic ductal adenocarcinoma cells^211^ suggesting that the disruption of SLIT2-ROBO1 signaling in pancreatic ductal adenocarcinoma may enhance metastasis via perineural invasion. Therefore, expression of the receptors *ROBO1* and *ROBO2* and two of their ligands *SLIT2* and *SLIT3* were characterized in a large number of pancreatic cancer cell lines and immortalized human pancreatic ductal epithelial cells (HPDE). To examine axon guidance molecule expression, total RNA was isolated, reverse transcribed, and queried with primers designed to span the 27^th^ intron of *ROBO1*, the 1^st^ and 15^th^ introns of *ROBO2*, the 33^rd^ intron of *SLIT2*, and the 35^th^ intron of *SLIT3*. As each of the axon guidance genes are very large, primers were designed as close to the 3’ end of each gene as possible to measure expression of the complete, or close to complete, transcript. qPCR analysis indicated that pancreatic cancer cell lines BxPC-3 and MIA PaCa-2 express approximately half the amount of *ROBO1* mRNA found in immortalized HPDE while HPAF-II and Hs 766T showed no detectable *ROBO1* expression. Su86.86 express an amount of *ROBO1* equal to HPDE while Capan-1 showed a 1.6-fold increase in *ROBO1* expression over HPDE, Capan-2 showed a 2.6-fold increase in *ROBO1* expression over HPDE, and PANC-1 showed a 6.2-fold increase in *ROBO1* expression over HPDE (Figure 2.1A). qPCR examination of *SLIT2* mRNA expression showed that pancreatic cancer cell line BxPC-3 express approximately half the amount of *SLIT2* found in HPDE while Su.86.86 express an amount of *SLIT2* equal to HPDE. Capan-1, Capan-2, HPAF-II, MIA PaCa-2, PANC-1, and Hs 766T do not express detectable levels of *SLIT2* (Figure 2.1B). qPCR analysis of *SLIT3* mRNA expression indicated that PANC-1 expressed *SLIT3* 515-fold higher than HPDE while the remaining pancreatic cancer cell lines examined showed little to no *SLIT3* expression (Figure 2.1C). qPCR of *ROBO2* mRNA using several different primer sets revealed that none of the pancreatic cancer cell lines used in this dissertation express *ROBO2* (data not shown).

**Figure 2.1:**
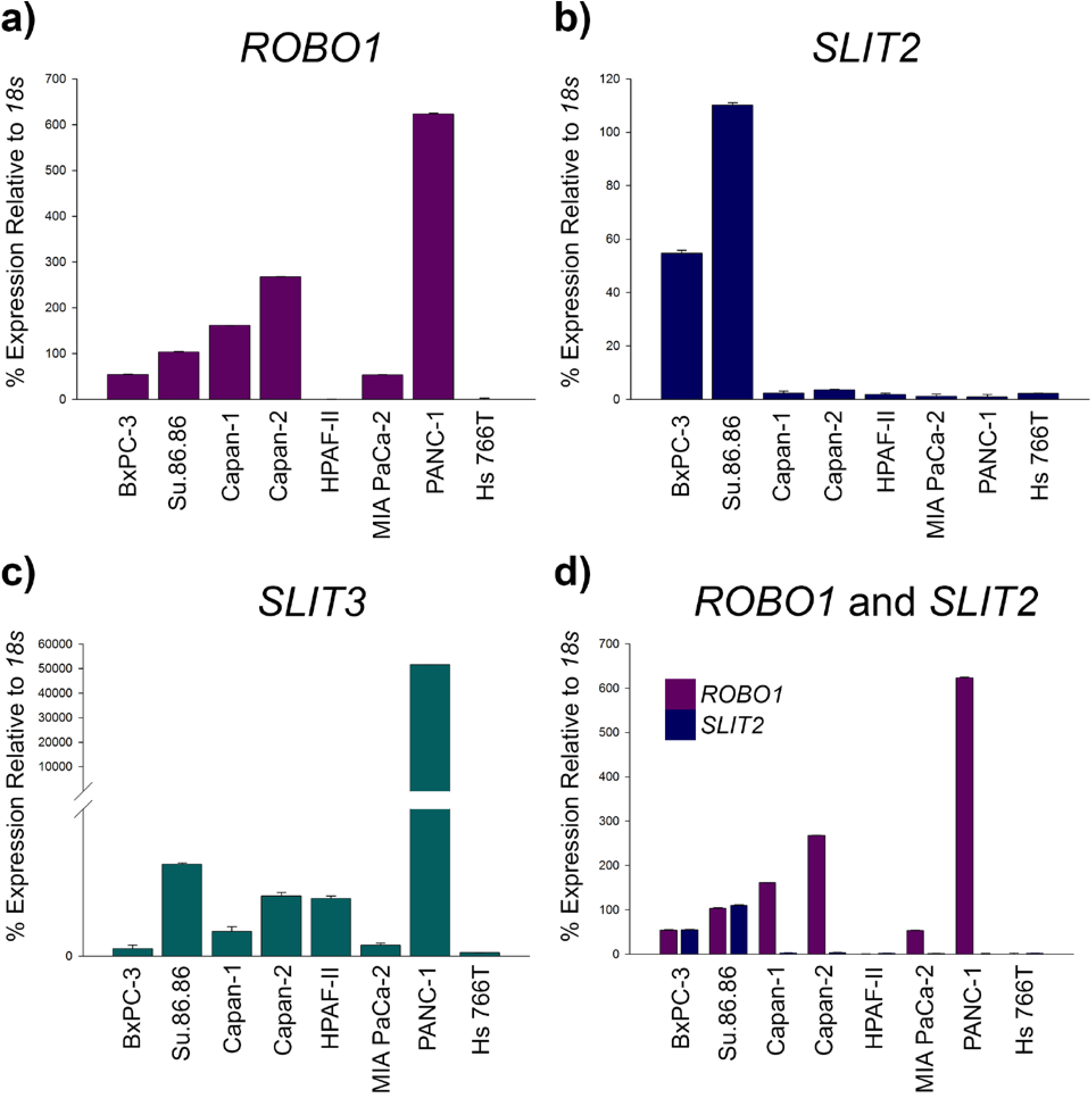
*ROBO1, SLIT2*, and *SLIT3* expression in pancreatic ductal adenocarcinoma cell lines. Total RNA was extracted with Trizol and cDNA was prepared using random primers and Superscript II. Quantitative PCR (qPCR) was carried out and mRNA levels were normalized to immortalized human pancreatic ductal epithelium relative to *18s*. Expression of mRNA as detected by qPCR for (a) *ROBO1*, (b) *SLIT2*, (c) *SLIT3*, and (d) *ROBO1* and *SLIT2. ROBO1* expression increases as pancreatic ductal adenocarcinoma cells gain independence from KRAS for growth and survival while *SLIT2* expression is lost in pancreatic ductal adenocarcinoma cells that contain activated KRAS. *SLIT3* expression is low in all pancreatic ductal adenocarcinoma cells examined except PANC-1. Pancreatic cancer cell lines can be classified into three subgroups based on their expression of *ROBO1* and *SLIT2*.

Cells depend on signaling through growth factors and their receptors to initiate signaling cascades that result in cell growth, proliferation and survival. Recently developed mouse models in which the Kras gene can be turned on and off have shown that oncogenic Kras signaling is necessary for the progression and maintenance of pancreatic ductal adenocarcinoma^235,236^. Sustained oncogeninc Kras signaling is also required for the growth and maintenance of metastatic pancreatic cancer lesions^237^. Pancreatic cancer cell lines with KRAS mutations contain elevated levels of KRAS activity^238^ and can either be dependent on KRAS signaling for growth and survival or independent of KRAS signaling for growth and survival^239^. As pancreatic ductal adenocarcinoma cancer cell lines gain independence from KRAS for growth and survival, *ROBO1* mRNA expression increases while *SLIT2* gene expression is lost in pancreatic cancer cell lines containing activated KRAS.

To assess protein expression and localization of ROBO1 and SLIT2, serial sections from cases of primary pancreatic tumors and distant normal were stained with antibodies to ROBO1 and SLIT2. In normal pancreas, ROBO1 is absent in the acinar cell compartment, ductal cells and the stroma (Figure 2.2A). In pancreatic ductal adenocarcinoma, only the ductal compartment shows strong ROBO1 staining (Figure 2.2B). In normal pancreas, SLIT2 is expressed strongly in both the acinar and ductal compartments while it remains absent in the stroma (Figure 2.2C) though not all sections contain SLIT2 staining in the acinar cell compartment. In pancreatic ductal adenocarcinoma, the ductal compartment shows weakened SLIT2 staining while it is absent in both the acinar cell compartment and stroma (Figure 2.2D). Reduction of SLIT2 expression in the ductal compartment of pancreatic ductal adenocarcinoma tissue sections along with the concurrent gain of ROBO1 expression in those same cells suggest that SLIT2/ROBO1 signaling is deregulated in pancreatic ductal adenocarcinoma.

**Figure 2.2:**
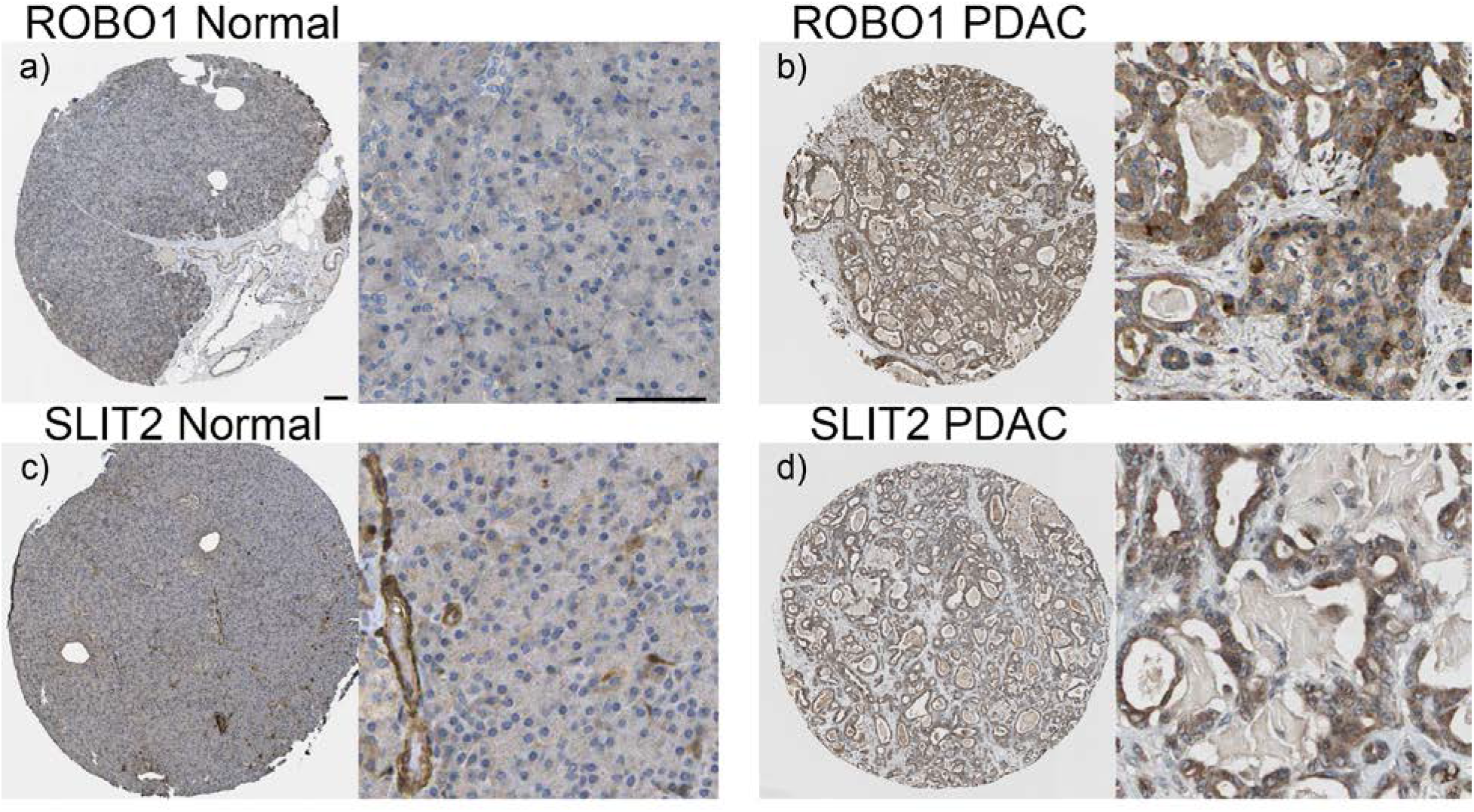
ROBO1 and SLIT2 expression in primary pancreatic ductal adenocarcinoma and distant normal. Deidentified, formalin-fixed, paraffin-embedded tissue was obtained from the University of Arizona GI SPORE tissue bank from patients that had not undergone preoperative chemotherapy or radiation therapy. Tissue sections were deparaffinized and immunohistochemistry was performed using either a rabbit polyclonal antibody to ROBO1 on (a) distant normal and (b) paired tumor or a rabbit polyclonal antibody to SLIT2 on (c) distant normal and (d) paired tumor. Staining for ROBO1 was absent in all cellular compartments in distant normal tissue sections while it showed strong staining in the ductal compartment in primary pancreatic ductal adenocarcinoma tissue sections. SLIT2 showed strong staining in both the acinar and ductal compartments in distant normal tissue sections, though not all tissue sections showed staining in the acinar compartment. Staining for SLIT2 showed weak staining in the ductal compartment in primary pancreatic ductal adenocarcinoma tissue sections. No tissue sections showed ROBO1 or SLIT2 staining in the stroma. Bar = 60 μm (a, left) and 100 μm (a, right). SLIT2/ROBO1 signaling is deregulated in pancreatic ductal adenocarcinoma.

### Epithelial and mesenchymal characteristics in pancreatic cancer cell lines

To become invasive and metastatic, epithelial cells undergo a process termed epithelial-to-mesenchymal transition which is characterized by the loss of epithelial cell characteristics such as cell polarity, organized structure, cell-cell contacts and expression of E-cadherin (*CDH1*) and the gain of mesenchymal cell characteristics including expression of transcriptional repressors *ZEB1*, Snail, and Slug^240^. Although SLIT2 expression is not restricted to epithelial cells—it can also be found in neurons and myoepithelial cells—to determine whether or not dysregulation of *ROBO1* and *SLIT2* in pancreatic cancer cells was associated with epithelial-to-mesenchymal transition, *CDH1* and *ZEB1* expression was examined in the pancreatic cancer cell lines.

Human pancreatic cancer cell lines BxPC-3, Su.86.86, Capan-1, Capan-2, and HPAF-II display an epithelial cell-like phenotype with an organized structure and defined cell-cell contacts. MIA PaCa-2, PANC-1, and Hs 766T cells; however, display a more mesenchymal cell-like phenotype with a more disorganized structure, a spindle-like morphology in MIA PaCa-2 and Hs 766T, and do not display intact cell-cell contacts. qPCR analysis of *CDH1* showed that BxPC-3, Su.86.86, Capan-1, Capan-2, and HPAF-II express epithelial cell marker *CDH1*, but do not express mesenchymal marker *ZEB1* further solidifying their epithelial phenotype (Figures 2.3A and B, respectively). MIA PaCa-2, PANC-1, and Hs 766T; however, do not express any *CDH1* mRNA, but do express very high levels of *ZEB1* further solidifying their mesenchymal phenotype (Figures 2.3A and B, respectively). Interestingly, gene expression of *ROBO1* or *SLIT2* does not correlate with expression of *CDH1* or *ZEB1*. These results suggest that dysregulation of *ROBO1* or *SLIT2* does not occur during the gain of invasive capabilities during epithelial-to-mesenchymal transition.

**Figure 2.3:**
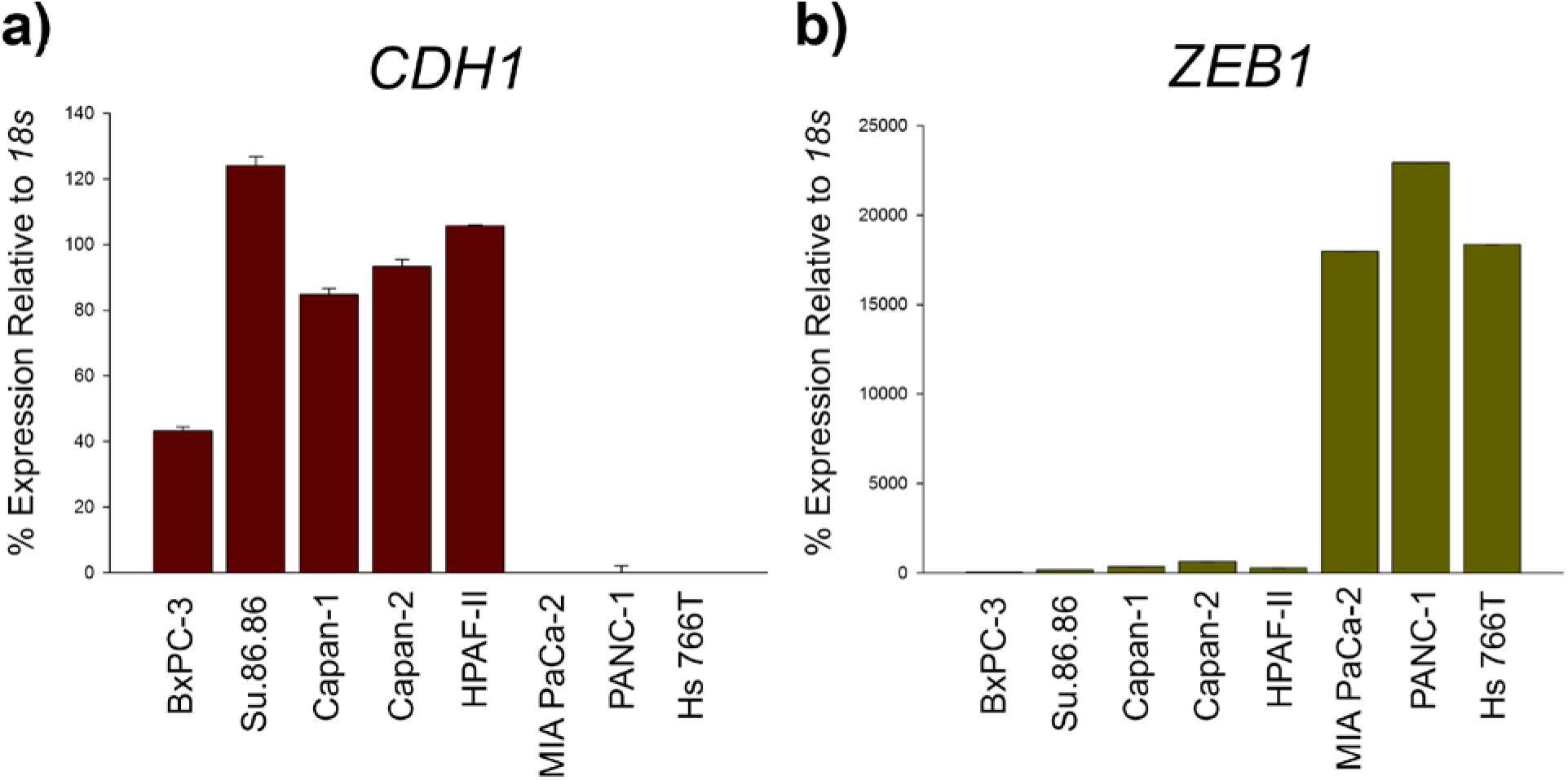
*CDH1* and *ZEB1* expression in pancreatic ductal adenocarcinoma cell lines. Total RNA was extracted with Trizol and cDNA was prepared using random primers and Superscript II. Quantitative PCR (qPCR) was carried out and mRNA levels were normalized to immortalized human pancreatic ductal epithelium relative to *18s*. Expression of mRNA as detected by qPCR for (a) *CDH1* and (b) *ZEB1*. Epithelial-like pancreatic ductal adenocarcinoma cell lines express *CDH1* while mesenchymal-like pancreatic ductal adenocarcinoma cell lines express *ZEB1*.

### DNA methylation of the SLIT2 core promoter

Expression of *SLIT2* has been shown to be silenced by methylation of its core promoter region in several other epithelial cell cancers^205-207^. In the two studies performed by Dallol et al. the *SLIT2* core promoter was defined as −761 bp to −212 bp upstream of the *SLIT2* transcriptional start site. Jin et al. analyzed the *SLIT2* promoter from −556 bp to −137 bp upstream of the transcriptional start site. All three studies used bisulfite sequencing to determine the methylation status of the *SLIT2* promoter. In the past, the gold standard for determining and quantifying the methylation state of DNA was by sequence analysis after bisulfite conversion (bisulfite sequencing)^241^; however, the cost of clonal sequencing can be costly and time consuming depending on the heterogeneity of DNA methylation. Recently, a novel approach for high-throughput DNA CpG methylation analysis has been introduced^242,243^. This method is based on a base-specific cleavage reaction combined with mass spectrometric analysis. This assay provides fast, quantitative screening of detailed methylation patterns using automated procedures that provides comparable results to bisulfite sequencing and also happens to be less time consuming and more cost-effective^244^. Therefore, Sequenom MassARRAY was utilized to examine the methylation status of the *SLIT2* core promoter in pancreatic cancer cell lines. Our analysis of the *SLIT2* promoter was divided into two overlapping primer sets that covered −611 bp to −79 bp upstream of the *SLIT2* transcriptional start site where R1 covers −344 to −79 bp and R2 covers −611 to −317 bp upstream of the *SLIT2* transcriptional start site. Sequenom analysis determined that the *SLIT2* core promoter in immortalized HPDE was almost completely unmethylated while various CpG fragments were methylated in all pancreatic cancer cell lines examined (Figure 2.4). In several gene promoters, a single or a few individual CpG sites often regulate complete transcriptional repression of gene expression^245–248^. Interestingly, the methylation status of fragment CpG:30 in Region 2 (−347 bp upstream of the *SLIT2* transcriptional start site) correlates with *SLIT2* expression in pancreatic cancer cell lines (Figures 2.1B and 2.4). The percent of methylation at CpG:30 in HPDE was 4.75% while the percent of methylation in BxPC-3 was 59%, Su.86.86 was 37.25%, Capan-1 was 98%, Capan-2 was 93.75%, and PANC-1 was 93% (Table 2.3). When compared to *SLIT2* expression, it is evident that the two pancreatic cancer cell lines with less than 60% methylation at CpG:30 express *SLIT2* while the three pancreatic cancer cell lines with greater than 60% methylation at CpG:30 do not. In addition, Su.86.86 which contains less than 40% methylation at CpG:30, shows expression of *SLIT2* at the same level as HPDE which has a methylation percentage of 4.75% at CpG:30.

**Figure 2.4:**
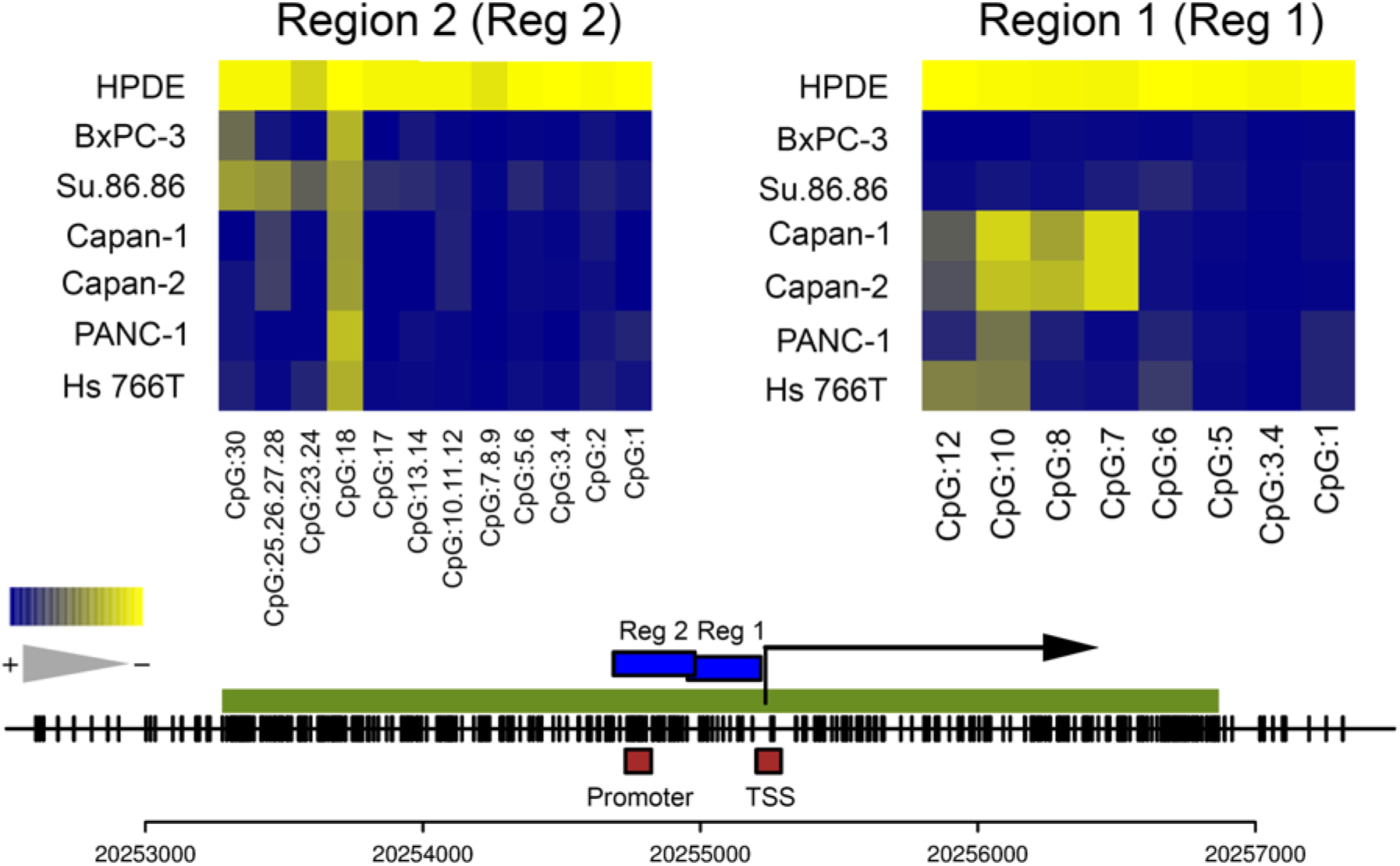
DNA CpG methylation within the *SLIT2* core promoter. Genomic DNA was isolated from pancreatic ductal adenocarcinoma cell lines and sodium bisulfite converted. Sodium bisulfite converted DNA was then amplified with region-specific PCR primers, *in vitro* transcribed, cleaved with RNaseA, and analyzed using a matrix-assisted laser desorption/ionization-time-of-flight mass spectrometer. Data is presented as the average percent methylation within each fragment. Yellow indicates 0% DNA CpG methylation while blue indicates 100% methylation. The green bar represents the large CpG island located within the *SLIT2* promoter, the blue bars represent the regions of the *SLIT2* promoter analyzed by Sequenom, the red boxes represent the regions analyzed by ChIP, and the numbers along the bottom represent the location in the genome in relation to the UCSC Genome browser hg19 assembly. The *SLIT2* core promoter remains unmethylated in immortalized human pancreatic ductal epithelium while it is methylated in all pancreatic ductal adenocarcinoma cell lines analyzed.

**Table 2.3:** Average percent methylation of fragment CpG:30 in the *SLIT2* core promoter. Genomic DNA was isolated from pancreatic ductal adenocarcinoma cell lines and sodium bisulfite converted. Sodium bisulfite converted DNA was then amplified with region-specific PCR primers, *in vitro* transcribed, cleaved with RNaseA, and analyzed using a matrix-assisted laser desorption/ionization-time-of-flight mass spectrometer. Data is presented as the average percent methylation for fragment CpG:30. Percent methylation of fragment CpG:30 correlates with *SLIT2* mRNA expression.

| Cell Line | Average % Methylation of Fragment CpG:30 |
| --- | --- |
| HPDE | 4.75 |
| BxPC-3 | 59.00 |
| Su.86.86 | 37.25 |
| Capan-1 | 98.00 |
| Capan-2 | 93.75 |
| PANC-1 | 93.00 |

To determine if DNA methylation of the *SLIT2* promoter or loss of *SLIT2* mRNA expression was a result of aberrant expression of DNA methyltransferases, expression of *DNMT1, DNMT3a*, and *DNMT3b* was characterized in our panel of pancreatic cancer cell lines. qPCR analysis indicated that pancreatic cancer cell lines BxPC-3, Su.86.86, Capan-2, HPAF-II, and PANC-1 express an amount of *DNMT1* mRNA equal to immortalized HPDE while Capan-1 and Hs 766T express approximately half the amount of *DNMT1* found in HPDE. MIA PaCa-2 showed a 3-fold increase in *DNMT1* expression over HPDE (Figure 2.5A). qPCR examination showed that BxPC-3 and Hs 766T express an amount of *DNMT3a* mRNA equal to HPDE while Su.86.86 and Capan-2 showed a 2fold increase in *DNMT3a* expression over HPDE, Capan-1 and PANC-1 showed a 3-fold increase in *DNMT3a* expression over HPDE, and HPAF-II and MIA PaCa-2 showed a 5.5 to 6-fold increase in *DNMT3a* expression over HPDE (Figure 2.5B). qPCR analysis indicated that BxPC-3, Su.86.86, and MIA PaCa-2 express approximately half the amount of *DNMT3b* found in HPDE whereas PANC-1 express an amount of *DNMT3b* mRNA equal to HPDE. Capan-2 and Capan-1 showed a 1.5 and 2-fold increase in *DNMT3b* expression over HPDE, respectively, while HPAF-II and Hs 766T showed a 4 to 4.5-fold increase in *DNMT3b* expression over HPDE (Figure 2.5C). Expression of *SLIT2* mRNA or DNA CpG methylation of the *SLIT2* promoter in pancreatic cancer cell lines does not correlate either directly or inversely with *DNMT1, DNMT3a*, or *DNMT3b* mRNA expression (Figure 2.1B, 2.4, and 2.5).

**Figure 2.5:**
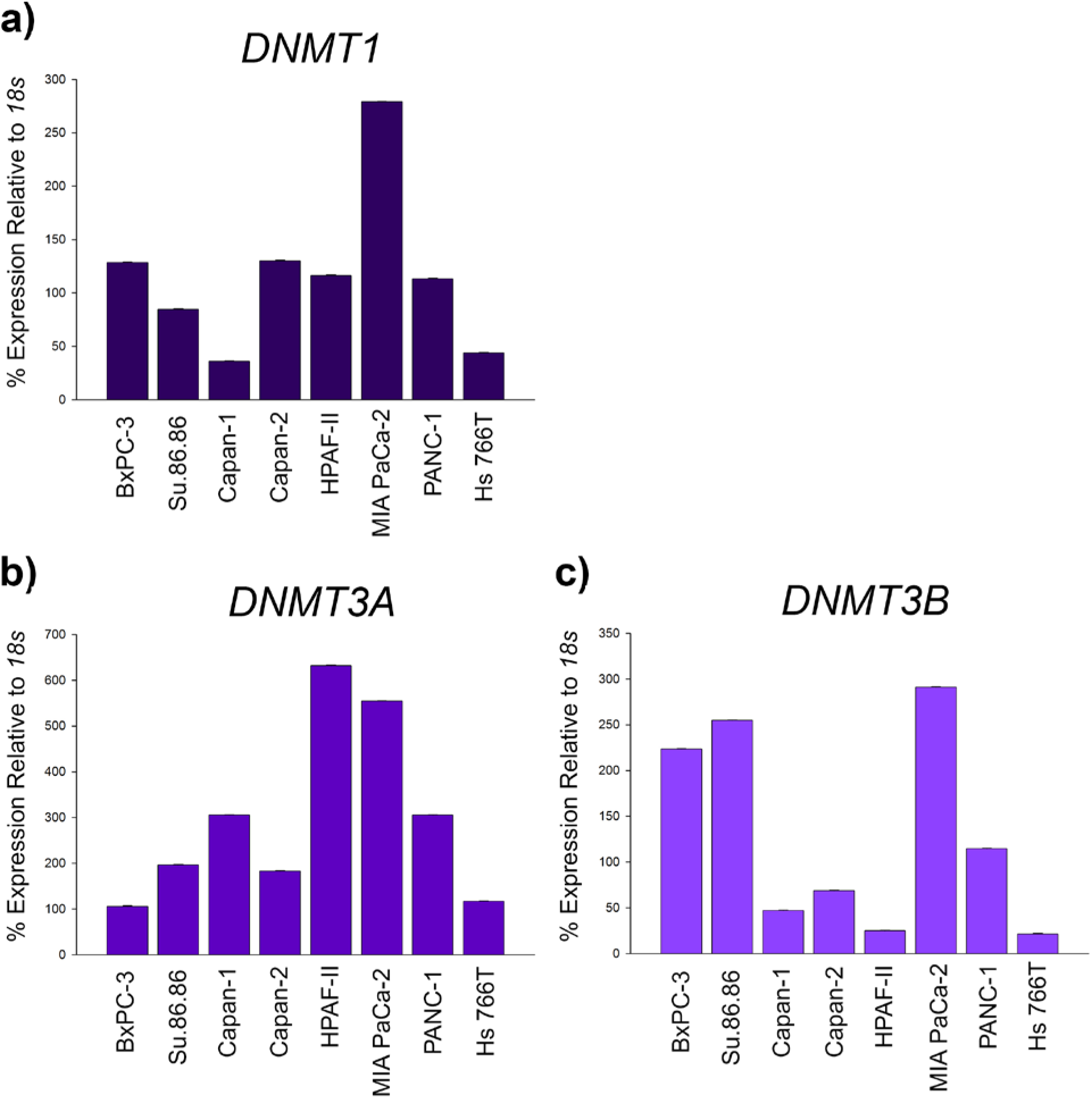
*DNMT1, DNMT3A*, and *DNMT3B* expression in pancreatic ductal adenocarcinoma cell lines. Total RNA was extracted with Trizol and cDNA was prepared using random primers and Superscript II. Quantitative PCR (qPCR) was carried out and mRNA levels were normalized to immortalized human pancreatic ductal epithelium relative to *18s*. Expression of mRNA as detected by qPCR for (a) *DNMT1*, (b) *DNMT3A*, and (c) *DNMT3B*. Expression of all three DNA methyltransferases is variable in pancreatic ductal adenocarcinoma cells and does not correlate with *SLIT2* mRNA expression or DNA CpG methylation of the *SLIT2* promoter.

Treatment with DNA demethylating agent 5-aza-2’deoxycytidine (5AdC) has been shown to cause re-expression of *SLIT2* in breast, lung, colon, and hepatocellular carcinoma cell lines^205–207^. Therefore, we treated the pancreatic cancer cell lines queried by Sequenom with 100 μM 5AdC to see if *SLIT2* can be re-expressed in these pancreatic cancer cell lines. In all pancreatic cancer cell lines treated with 5AdC, *SLIT2* expression was reactivated (Figure 2.6). Histone deacetylase inhibitors can also increase gene expression in the absence of hypermethylation^49^. Since DNA methylation is dominant to histone deacetylation, inhibition of methylation must occur before transcription can. Therefore, treatment with demethylating agents such as 5AdC followed by histone deacetylase inhibitors like Trichostatin A (TSA) can produce an additive or synergistic effect for gene re-expression^249,250^. Treatment with 50 nM TSA alone did not cause *SLIT2* re-expression in any pancreatic cancer cell line, and treatment with both 5AdC and TSA did not result in an additive or synergistic effect for *SLIT2* re-expression (Figure 2.6). These results suggest that the *SLIT2* core promoter is methylated in pancreatic cancer, the percentage of methylation at a single CpG site correlates with *SLIT2* expression, and *SLIT2* mRNA can be re-expressed after treatment with demethylating agent 5AdC, but not histone deacetylase inhibitor TSA or a combination of the two drugs.

**Figure 2.6:**
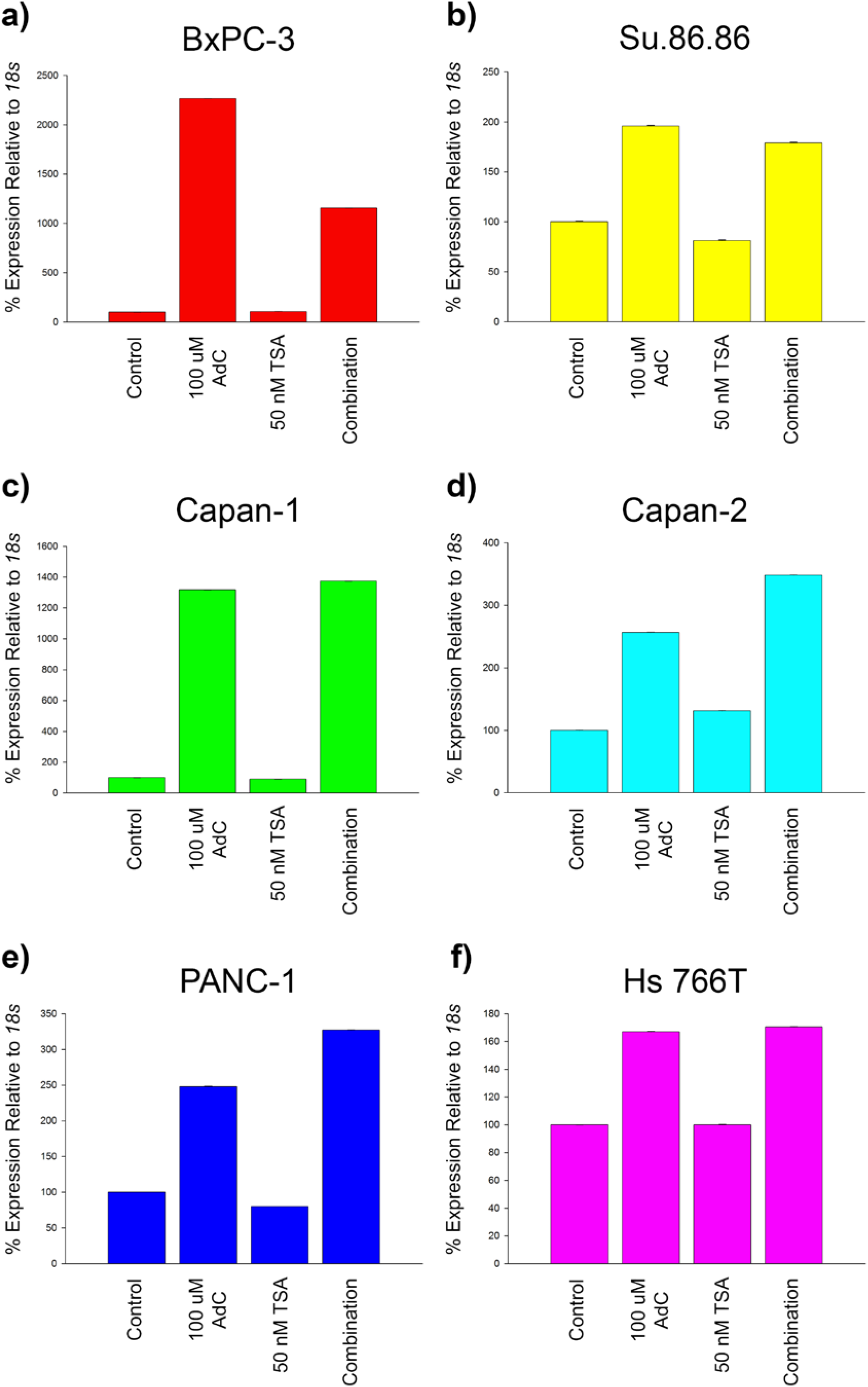
*SLIT2* re-expression in pancreatic ductal adenocarcinoma cells. Cells were treated for six days with 100 μM 5-aza-2’deoxycytidine (5AdC) or for twenty-four hours with 50 nM Trichostatin A (TSA). Cells treated with a combination of the two drugs were treated for six days with 100 μM 5AdC and 50nM of TSA added twenty-four hours before collection. Total RNA was extracted with Trizol and cDNA was prepared using random primers and Superscript II. Quantitative PCR (qPCR) was carried out and mRNA levels were normalized to control non-treated cells relative to *18s*. Re-expression of *SLIT2* mRNA as detected by qPCR in (a) BxPC-3, (b) Su.86.86, (c) Capan-1, (d) Capan-2, (e) PANC-1, and (f) Hs 766T. *SLIT2* mRNA can be re-expressed after treatment with demethylating agent 5AdC, but not histone deacetylase TSA or a combination of the two drugs.

### Histone modifications at the SLIT2 promoter and transcriptional start site

Expression of *SLIT2* has also been shown to be negatively regulated by H3K27 trimethylation (H3K27me3) by the PRC2 component EZH2. Hence, ChIP was performed to map out which histone modifications are present in the *SLIT2* promoter and at the *SLIT2* transcriptional start site in pancreatic cancer cells. ChIP primers to the *SLIT2* promoter were designed from −568 bp to −474 bp upstream of the *SLIT2* transcriptional start site while ChIP primers to the *SLIT2* transcriptional start site were designed from −96 bp to −6 bp upstream of the *SLIT2* transcriptional start site. Dimethylation of H3K4 (H3K4me2) and acetylation of H4 (H4ac) were used as markers of potential *SLIT2* transcriptional activation while H3K27me3 was used as a marker of potential *SLIT2* transcriptional repression. To look at the role of these histone modifications and compare them to *SLIT2* expression we used immortalized HPDE, Su86.86 since they express *SLIT2* transcripts, and PANC-1 because they show no *SLIT2* mRNA expression. In the *SLIT2* promoter region (ChIP primer encompassing −568 bp to – 474 bp and correlating to the first half of Region 2 analyzed by Sequenom MassARRAY), chromatin enrichment of H4ac in HPDE was 6.77-fold over the no antibody control, Su.86.86 was 15.24-fold, and PANC-1 was 5.17-fold. Chromatin enrichment of H3K4me2 in HPDE was 14.12-fold, Su.86.86 was 13.27-fold, and PANC-1 was 3.66-fold. Enrichment of H3K27me3 in HPDE was 0.50-fold, Su.86.86 was 1.72-fold, and PANC-1 was 1.87-fold. Overall, HPDE showed moderate chromatin enrichment for activating histone mark H3K4me2, Su.86.86 contained moderate chromatin enrichment for activating histone marks H3K4me2 and H4ac, and both cell lines showed low-to-no chromatin enrichment of the inactivating histone mark H3K27me3 in the *SLIT2* promoter (Figure 2.7A). At the *SLIT2* transcriptional start site (ChIP primer encompassing −96 bp to −6 bp and located downstream of Region 1 analyzed by Sequenom MassARRAY), enrichment of H4ac in HPDE was 4.26-fold over the no antibody control, Su.86.86 was 11.35-fold, and PANC-1 was 2.20-fold. Enrichment of H3K4me2 in HPDE was 8.00-fold, Su.86.86 was 11.59-fold, and PANC-1 was 1.52-fold. Enrichment of H3K27me3 in HPDE was 1.39-fold, Su.86.86 was 6.48-fold, and PANC-1 was 3.79-fold. Interestingly, performing ChIP with an antibody to RNA polymerase II showed that the *SLIT2* transcriptional start site was enriched at 2.33-fold over the no antibody control in HPDE, Su.86.86 was 2.45-fold, and PANC-1 was 1.43-fold. Overall, the two cell lines that express *SLIT2* mRNA, HPDE and Su.86.86, had higher chromatin enrichment of activating histone modifications H3K4me2 and H4ac at the *SLIT2* transcriptional start site as well as a higher enrichment for RNA polymerase II while PANC-1—which does not express *SLIT2* mRNA at detectable levels—has higher chromatin enrichment of inactivating histone mark H3K27me3 and a lower chromatin enrichment for RNA polymerase II (Figure 2.7B). Since EZH2 is the enzyme responsible for depositing methyl groups onto H3K27, we sought to measure *EZH2* mRNA levels in pancreatic cancer. Expression of *EZH2* mRNA in pancreatic cancer cell lines is inversely correlated with *SLIT2* mRNA expression (Figure 2.1B and 2.7C). Previous work in prostate cancer has shown that *SLIT2* is a target of EZH2 repression, its expression can be restored by treatment with demethylating agent 5AdC, histone deacetylase inhibitor SAHA, and EZH2 inhibitor 3-Deazaneplanocin A, and that the addition of SLIT2 inhibited prostate cancer proliferation and invasion^208^. These results suggest that *SLIT2* gene expression is controlled by histone modifications at the *SLIT2* transcriptional start site, but not the *SLIT2* promoter.

**Figure 2.7:**
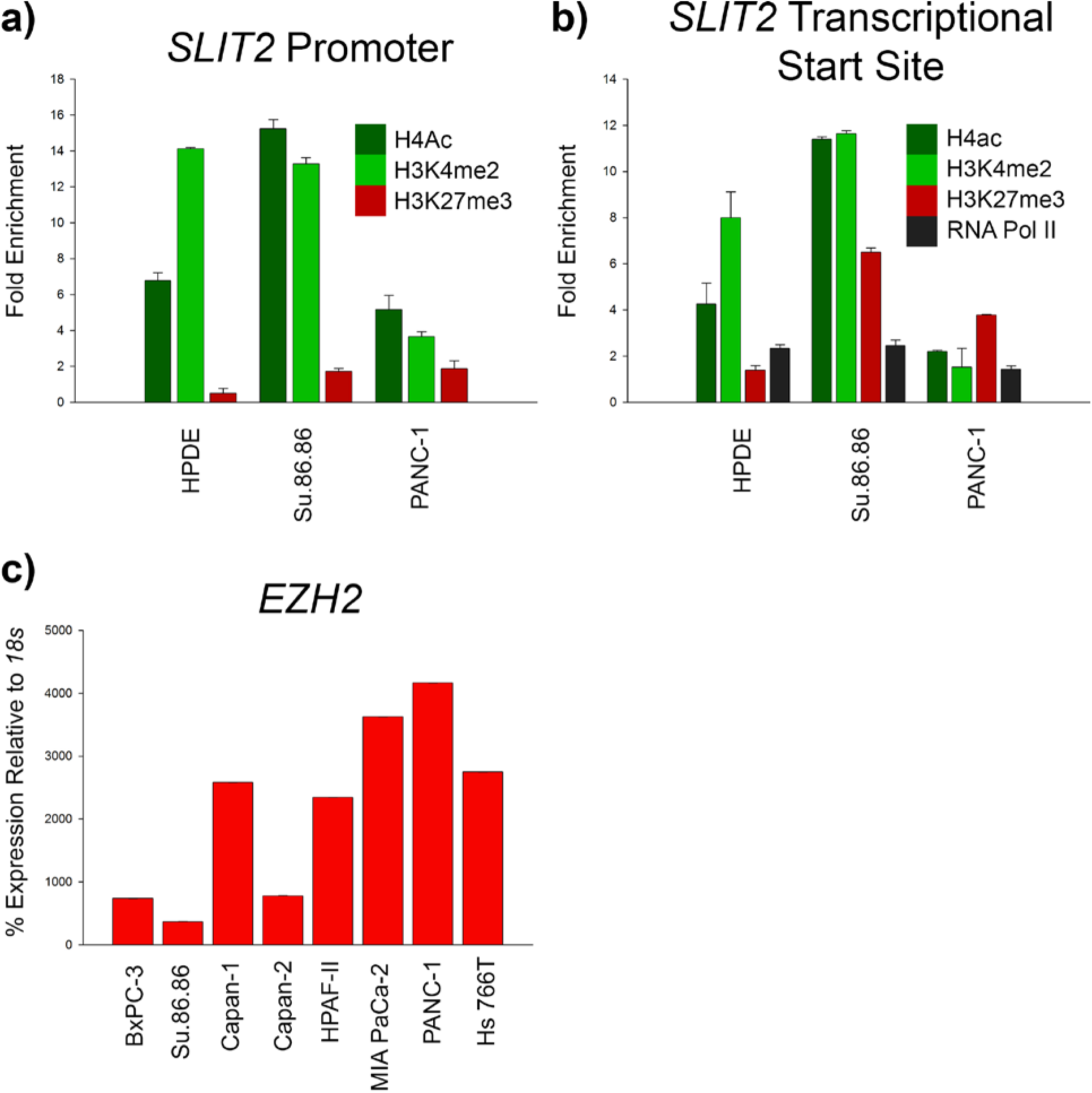
Chromatin enrichment of histone modifications at the *SLIT2* promoter and transcriptional start site and *EZH2* expression in pancreatic ductal adenocarcinoma cells. Cell lysates were crosslinked, scraped, sonicated, precleared, and incubated overnight with rabbit polyclonal antibodies to H4ac, H3K4me2, and H3K27me3. Lysates were also incubated overnight with a mouse monoclonal antibody to RNA Polymerase II. 10% of cell lysates were set aside as input DNA prior to preclearing. Immunoprecipitated DNA was washed, crosslinks were reversed, protein was digested, and equal amounts of immunoprecipitated DNA and input DNA were analyzed via quantitative PCR (qPCR). A difference in immunoprecipitation was determined using the fold enrichment method. (a) Chromatin enrichment for H4ac, H3K4me2, and HeK27me3 at the *SLIT2* promoter. (b) Chromatin enrichment for H4ac, H3K4me2, H3K27me3, and RNA Polymerase II at the *SLIT2* transcriptional start site. Total RNA was extracted with Trizol and cDNA was prepared using random primers and Superscript II. qPCR was carried out and mRNA levels were normalized to human pancreatic ductal epithelium relative to *18s*. (c) Expression of *EZH2* mRNA as detected by qPCR. HPDE and Su.86.86 show higher chromatin enrichment of activating histone modifications H3K4me2 and H4ac at the *SLIT2* transcriptional start site as well as a higher enrichment for RNA polymerase II while PANC-1 show higher chromatin enrichment of inactivating histone mark H3K27me3 and a lower chromatin enrichment for RNA polymerase II. Expression of *EZH2* inversely correlates with *SLIT2* mRNA expression.

## Discussion

In the present study we have evaluated the gene expression of axon guidance receptor *ROBO1* and its ligand *SLIT2* in pancreatic ductal adenocarcinoma. We found that in normal pancreas, high levels of SLIT2 are seen in the ductal compartment of every patient and in the acinar cell compartment of the occasional patient whereas ROBO1 antibody staining is not seen within any pancreatic cell compartment. In primary human pancreatic ductal adenocarcinomas, SLIT2 staining is absent in the acinar cell compartment and decreased, but not completely absent, in the ductal cells. ROBO1 staining; however, is strong in the ductal compartment. The same pattern of expression can be seen at the RNA level in pancreatic cancer cell lines. As pancreatic cancer cell lines become KRAS-independent^238^, *ROBO1* expression increases while *SLIT2* expression is lost in pancreatic cancer cell lines that contain activated Kras. Only one pancreatic cancer cell line expresses *SLIT3*. Interestingly, we saw that pancreatic cancer cell lines fall into three distinct groups based on *ROBO1* and *SLIT2* expression: 1) cells that express both *ROBO1* and *SLIT2* such as HPDE, BxPC-3, and Su.86.86; 2) cells that express only *ROBO1* such as Capan-1, Capan-2, MIA PaCa-2, and PANC-1; and 3) cells that express neither *ROBO1* nor *SLIT2* such as HPAF-II and Hs 766T (Figure 2.1D). Upon comparison to expression of *CDH1* and *ZEB1*, we can conclude that expression of *ROBO1* and *SLIT2* does not correlate with gene expression changes associated with epithelial-to-mesenchymal transition. It has long been known that breast cancers can be classified into four main molecular subtypes based on gene expression patterns^251^ and correlates with clinical outcome. The clear separation of pancreatic cancer cell lines based on *ROBO1* and *SLIT2* expression suggest that specific subtypes of pancreatic cancers exist and may be defined by their expression of these two axon guidance molecules and that *ROBO1* or *SLIT2* expression could be used to determine clinical outcome and survival in patients with pancreatic ductal adenocarcinoma.

Gene expression can be positively or negatively regulated. One of the most ubiquitously used mechanisms of gene repression a cell uses to silence tumor suppressor genes is DNA methylation of CpG sites within a gene promoter. Epigenetic silencing through this mechanism occurs as frequently as mutations or deletions^60,61^. It has already been shown that the *SLIT2* promoter is heavily methylated in several other types of epithelial cancers^205–207^. Vincent et al. (2011) found that SLIT2 was hypermethylated in pancreatic cancer cells, but remained unmethylated in immortalized HPDE cells^252^. In a more recent study, Nones et al. took samples from nontreated primary pancreatic ductal adenocarcinoma patients and adjacent normal to determine differentially methylated regions in pancreatic cancer. They found that *SLIT2* was methylated in the pancreatic cancer samples, but not in the adjacent normal which was then validated by deep-sequencing of bisulfite converted DNA. This group also noticed an inverse correlation between *SLIT2* methylation and gene expression^253^ suggesting epigenetic inactivation plays a role in the disruption of SLIT-ROBO signaling in pancreatic ductal adenocarcinoma. Here we used Sequenom MassARRAY technology to confirm DNA methylation of the *SLIT2* promoter. We found that the *SLIT2* promoter remains unmethylated in immortalized HPDE while there are varying levels of promoter methylation in pancreatic cancer cells with promoter methylation increasing alongside the activation of Kras. What is interesting; however, is that even though Su.86.86, which contain activated Kras, have approximately 38% promoter methylation, they express *SLIT2* at a level equivalent to HPDE. This may suggest a threshold of methylation that must be met in order for complete gene silencing, or that specific CpG sites within the *SLIT2* promoter must be methylated for complete gene silencing to occur. In this study, methylation of CpG:30 in Region 2 correlates with our *SLIT2* expression data. Next to HPDE, Su.86.86 has the least amount of overall DNA methylation in the *SLIT2* core promoter and the highest level of *SLIT2* expression out of all the pancreatic cancer cell lines analyzed. BxPC-3 has slightly more DNA methylation than Su.86.86 and slightly lower *SLIT2* expression than HPDE. Finally, Capan-1, Capan-2, PANC-1, and Hs 766T all have 85-100% DNA methylation at CpG:30 and contain no expression of *SLIT2*. Another possibility is that the region encompassing CpG:30 contains a transcription factor binding element that may attract chromatin remodeling complexes leading to the stimulation of transcription. However, definitive studies to map the key positive and negative promoter elements for *SLIT2* have not been performed.

DNA methyltransferases are the enzymes responsible for depositing methyl groups onto cytosines and have been shown to be expressed in primary pancreatic ductal adenocarcinoma tissue^254^. They are also responsible for regulating gene expression^255^. mRNA expression of *DNMT1, DNMT3a*, and *DNMT3b* was found to be variable in pancreatic cancer cell lines and did not correlate either directly or indirectly with *SLIT2* expression. DNA methylation is also reversible upon treatment with demethylating agents such as 5-aza-2’deoxycytidine (5AdC) including DNA methylation and re-expression of *SLIT2*^205-207^. In this study, every pancreatic cancer treated with 100 μM 5AdC induced *SLIT2* expression; however, the amount of induction depended on basal *SLIT2* expression. For example, pancreatic cancer cell lines with low basal *SLIT2* expression had a higher re-expression upon treatment. The presence of DNA methylation and its correlation with *SLIT2* expression suggest that it is a major component of *SLIT2* transcriptional repression in pancreatic ductal adenocarcinoma. DNA methylation of *SLIT2* could also potentially play a role in pancreatic cancer diagnosis or patient stratification. Re-expression of *SLIT2* upon treatment with 5AdC makes it a tantalizing drug target as well.

Chromatin structure also determines the expression pattern of genes. Histone acetylation is required to keep chromatin in an open state that is optimal for gene transcription^50^. Histone deacetylase (HDAC) proteins commonly bind to the promoter region of genes to cause histone deacetylation, and, therefore, a loss of gene expression^256^. Along with DNA methylation, histone deactylation is reversible upon treatment with histone deacetylase inhibitors. In this study, we treated pancreatic cancer cell lines with pan-histone deacetylase inhibitor Trichostatin A (TSA). Upon treatment with 50 nM TSA, *SLIT2* was not re-expressed though it was not confirmed whether treatment with TSA altered histone acetylation. It has long been known that DNA methylation is the dominant epigenetic mark and that gene re-expression cannot occur without first inhibiting methylation^249,250^. Co-treatment of pancreatic cancer cell lines with 100 μM 5AdC and 50 nM TSA did not lead to an additive or synergistic effect on *SLIT2* re-expression. Lack of *SLIT2* re-expression upon treatment with a pan-histone deacetylase inhibitor suggests that histone deactylation is not a mechanism of *SLIT2* gene expression in pancreatic ductal adenocarcinoma.

Specific histone modifications add an additional layer of complexity in gene regulation. Histone modifications can occur in different histone proteins (i.e. H3 and H4), different histone variants (i.e. H3.3 and H2A.Z), and different histone residues (i.e. lysine, arginine, and serine). The modifications may be methyl groups, acetyl groups, or phosphorylation and methylation can occur as mono, di, or trimethylation. Specific histone modifications are known to be either activating or repressive. Activating marks include acetylation of H3 and H4, H3K9 acetylation, and H3K4 di and trimethylation. Repressive marks include loss of histone acetylation, H3K9 methylation, and H3K27 trimethylation. Loss of activating histone marks and the gain of repressive marks often occurs alongside DNA methylation^60^. We utilized chromatin immunoprecipitation to analyze the chromatin landscape of the *SLIT2* promoter and transcriptional start site. We found that HPDE and Su86.86 both contain at least two-times the amount of chromatin enrichment for activating histone marks H4ac and H3K4me2 than inactivating histone mark H3K27me3 at the *SLIT2* transcriptional start site. PANC-1, however, have approximately twice the amount of chromatin enrichment for inactivating histone mark H3K27me3 than activating histone marks H4ac and H3K4me2. Since there is a higher ratio of chromatin enrichment for activating histone marks in both HPDE and Su.86.86, this suggests that the cellular environment around the *SLIT2* transcriptional start site is conducive to transcription. In PANC-1, there is a higher ratio of chromatin enrichment for inactivating histone mark H3K27me3 suggesting that the cellular background in these cells promotes transcriptional repression. While the amount of chromatin enrichment was never greater than 15-fold for any histone mark, and the majority of chromatin enrichment was less than 5-fold, this work suggests that the transcriptional start site of *SLIT2* has the potential of being a bivalent domain in pancreatic cancer cells capable of *SLIT2* transcription depending on the cellular context^228^.

In 2006, it was discovered that bivalent domains strongly correlate with CpG islands^229^. CpG islands also play an important role in establishing and maintaining H3K27me3 enrichment at bivalent domains^228^ with more localized patterns of H3K27me3 distribution around the transcriptional start site^230,257^. In this study, there was more chromatin enrichment for H3K27me3 at the *SLIT2* transcriptional start site than in the *SLIT2* promoter. Interestingly, defined peaks of components of the polycomb repressive complex 2—which contains the enzyme EZH2—are found around gene promoters^77,231,258^. In human embryonic stem cells, approximately 97% of all promoter-associated EZH2-recruitment sites correspond to CpG islands that contain binding sites for transcription factors that act in a repressive manner^231^. Conversely, the same study showed that EZH2-free CpG islands contain binding sites for transcription factors that promote transcription. Thus, CpG islands in promoter regions may maintain a bivalent conformation not only through histone modifications, but through transcription factors as well suggesting that bivalency acts as a means to fine-tune expression of genes. The presence of both activating and repressive histone marks at the *SLIT2* transcriptional start site suggests the potential presence of a bivalent domain. It is possible that the large CpG island that exists −2021 base pairs upstream of the *SLIT2* transcriptional start site and +1566 base pairs downstream of the transcriptional start site plays a role in establishing enrichment of H3K27me3 at this location in the genome. Considering that *EZH2* expression is also inversely correlated with *SLIT2* expression, expression of *SLIT2* may depend on the activity and function of *EZH2* outside of its ability to deposit methyl groups on H3K27.

This work suggests that loss of *SLIT2* expression may play a role in the progression of pancreatic ductal adenocarcinoma. It is likely that the loss of *SLIT2* in pancreatic cancers with activated Kras is due to epigenetic silencing through both DNA methylation and histone modifications (Figure 2.8). Since *SLIT2* has properties of a tumor suppressor gene (inhibition of proliferation, migration and invasion), this epigenetic silencing of *SLIT2* may facilitate tumor progression. Future directions would be to determine if there is a mechanistic link between KRAS activation and DNA methylation of the *SLIT2* promoter and to elucidate the transcription factors associated with *SLIT2* transcriptional activation and repression that bind to areas of the large, potentially EZH2-associated CpG island located in the *SLIT2* promoter.

**Figure 2.8:**
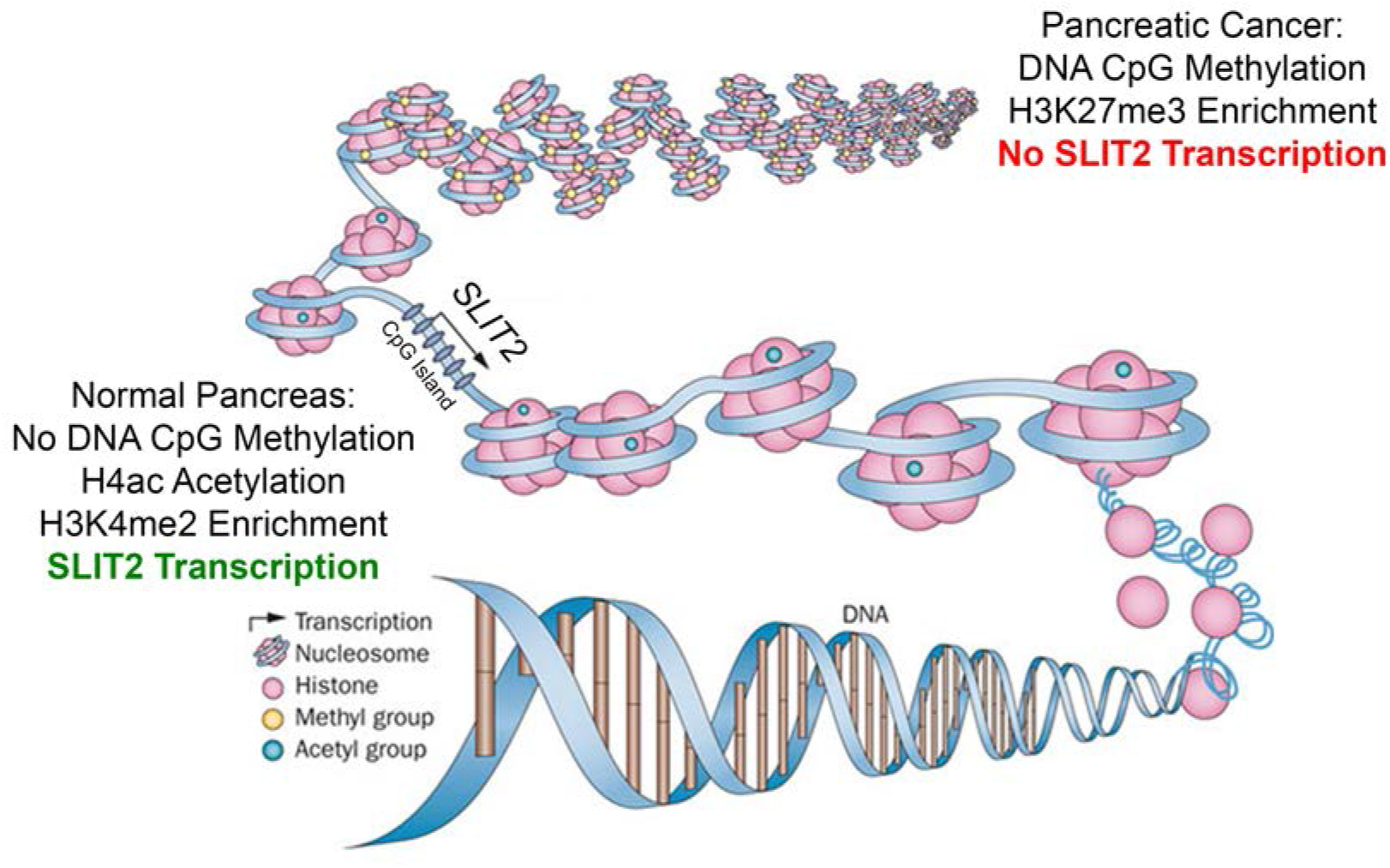
Loss of *SLIT2* is due to epigenetic silencing. In normal pancreas cells, the *SLIT2* promoter contains DNA with no CpG methylation within the CpG island and activating histone modifications such as H4ac and H3K4me2 around the *SLIT2* transcriptional start site allowing for transcription to occur. When pancreatic ductal adenocarcinomas gain independence from KRAS for growth and survival, the *SLIT2* promoter gains DNA CpG methylation within the CpG island and the repressive histone modification H3K27me3 at the *SLIT2* transcriptional start site blocking transcription. Schematic adapted from Azad et al. Nat Rev Clin Oncol. 2013 May; 10(5): 256-66.

This work is the first to show cell autonomous expression of the SLIT2/ROBO1 pathway in pancreatic ductal adenocarcinoma. Using *ROBO1* and *SLIT2* gene expression as a way to classify pancreatic ductal adenocarcinomas may prove useful in furthering the molecular characterization of this disease. This work is also the first to show potential bivalency of the *SLIT2* gene promoter. Furthermore, epigenetic regulation of *SLIT2* mRNA expression through DNA methylation of its CpG island has the potential for use as a biomarker in detecting pancreatic ductal adenocarcinomas—a helpful tool since no tests for early detection currently exist.

